# Characteristics of executive control network function based on a lateralized version of the ANT-R test in athletes from open skill sports: An fNIRS study

**DOI:** 10.1101/540500

**Authors:** Miao Yu, Yi B. Liu, Guang Yang

## Abstract

The purpose of the study was to investigate the executive control network function characteristics of interceptive and strategic sports athletes from open skill sports. In order to do so, we used a revised lateralized attention network task to measure executive control efficiency and activation related to flanker interference changes on the right frontoparietal network using functional near-infrared spectroscopy in athletes from different sport sub-categories. Strategic athletes had higher accuracy and lower flanker conflict effects on accuracy, as well as longer reaction time and stronger conflict effects under the valid cue and invalid cue conditions. This was accompanied by higher activity in the right inferior frontal gyrus. These results extend the evidence suggesting that differences among interceptive sports and strategic sports athletes are due to the former using higher velocities to solve conflicts, and the latter using higher accuracy in the same tasks. These effects are attributed to differences in the right frontoparietal network.

## Introduction

Growing evidence indicates that elite athletes are better able to integrate skills and tactics to achieve peak performance [1,2]. The fundamental difference between excellent athletes and novices or non-athletes is related to sophisticated and specific cognitive processing using the optimization of neurocognitive resources, rather than simple physical ability and skill. Jacobson and Matthaeus [3] found not only that athletes outperform non-athletes on tests of executive function domains such as inhibition and problem solving, but also that varying athletic experience may correlate with higher functioning of particular executive domains.

Sports may be categorized into two types: open skill and closed skill [3,4]. Previously, most studies have focused on establishing associations between chronic [5,6] or acute open and closed skill exercise [7], cognitive proficiency, and reorganization in the human brain [4], in addition to comparing the impact of open and closed skills on cognition [4,8,9]. Recent neuroimaging techniques have revealed that long-term motor skill training causes plastic reorganization of brain structure and function professional athletes playing various sports, including closed skill athletes (such as world class gymnasts [10,11], world class mountain climbers [12], and formula racing-car drivers [13]), and open skill athletes (such as badminton [14] and table tennis players [15]). The results of these studies suggests that brain reorganization may vary across different types of athlete.

In addition, open and closed-skill sports can be further subcategorized. Of note, sub-categories of open skill sports include interceptive (e.g., table tennis, fencing, and tennis) and strategic (e.g., sports that involve multiple teammates, opponents, and tactical formations, such as basketball, football, etc. [2,3,7]). Little is known about whether plastic changes can be modulated differentially across sport sub-categories as a result of the differences in cognitive and motor demands.

Executive control function is fundamentally important for success in open skill sports, because it manages other basic cognitive functions such as inhibition of behavior, attention, and working memory [19]. A study by Vestberg et al. [20] strongly suggested that results of executive function tests predict the success of ball sport players. To date, differences in the interplay between executive function and associated neural mechanisms in interceptive versus strategic sports remain to be elucidated. However, numerous previous studies have examined executive function by using variants of the color Stroop task and the flanker task [21].

Attention is an important aspect of executive function, serving as the basis for various neural regulatory systems. Attention is manifest in the activity of a set of brain networks that influence the priority of computations of other brain networks enabling appropriate access to consciousness and observable behavior [16,17]. The executive control function of attention involves complex mental operations in detecting and resolving conflict [18], and has also been used to evaluate the efficiency of executive control [21].

In competition, athletes’ executive control does not only involve planning actions, decision-making and overcoming habitual actions according the trajectories and falling positions of dynamic equipment, but also detection of feint actions by opposing players, judging the genuine intentions of opponents and determining interactions with dynamic cues (e.g. opponents’ body positions, facial expressions, and position of the ball or racket). These judgments are more closely related to movement as opposed to functions assessed by tasks, such as Flanker tests.

Clinical studies in patients with brain lesions have indicated right hemisphere dominance for the executive network is characteristic of most humans [22]. Further, a larger parieto-frontal network in the right than left hemisphere was significantly related to the degree of anatomical lateralization and asymmetry of performance on visuospatial tasks [25].

Therefore, we used a revised lateralized attention network test (LANT-R) to investigate the characteristics of executive control in athletes from different sport sub-categories including table tennis, tennis, badminton, football, basketball, and volleyball. We also compared athletes to average non-athlete college student controls. In this paper, we present data from a preliminary study investigating the relationships between sport type and the neural mechanisms of executive control, and propose a theory regarding the role of cues in executive control.

In contrast to other neuroimaging methods, multichannel functional near-infrared spectroscopy (fNIRS) is portable and uses compact experimental systems. Measurements obtained using fNIRS are limited to lateral cortical surfaces. Neuroimaging techniques have revealed that the executive control function is mainly associated with activation of the extended frontoparietal network (FPN) [24], with right hemisphere dominance for attention [22, 25–26]. Therefore, we investigated possible differences in efficiency of executive attention networks and activation related to flanker interference changes in the right FPN. To our knowledge, this study provides the first experimental evidence that differences exist in the neural mechanisms of executive control in athletes during interceptive versus strategic sports. We hypothesized that all athletes, regardless of sport sub-type, would demonstrate significantly faster reaction times (RT) and higher accuracy (ACC) when detecting and resolving conflicts, smaller flanker conflict effects, and higher activity in the right dorsal lateral prefrontal cortex (rdLPFC) than non-athletes. We further posited that there are significant differences in executive control function between interceptive and strategic sports athletes under different cue conditions, and that variability in performance was related to activation of the right inferior frontal gyrus (rIFG).

## Methods

### Participants

Fifty-six athletes participated in the study. Participants were recruited through poster advertisements at Jilin Sport University. The subjects comprised 16 interceptive sports athletes played table tennis (n = 6), badminton (n = 5), and tennis (n = 5), and 16 strategic sports athletes who played basketball (n = 6), volleyball (n = 5), and soccer (n = 5). The athletes satisfied all of the following criteria: (1) had 5 or more years of professional training experience, (2) qualified as a national player in the second grade or above, (3) trained more than five times per week in the last four years, and (4) trained for two or more hours at a time. The non-athlete control group comprised 24 students majoring in rehabilitation, who matched the athlete group in age and education, but had no experience in playing sports or athletic training. All participants were right-handed and had normal or corrected-to-normal visual acuity. No individuals reported having a history of neurological or psychiatric disorder. Table 1 shows the main characteristics of the subjects. All procedures were approved by the Ethics Committee of the Jilin Sport University. Informed written consent was obtained from each participant prior to the experiment, and they were paid an inconvenience allowance.

**Table 1.** Demographic and physical characteristics of the participants in each group

| Variables | strategic sports (n = 16) | interceptive sports (n = 16) | Control (n = 24) |
| --- | --- | --- | --- |
| Female | 8 | 9 | 12 |
| Age (years) | 22.13 (1.37) | 21.67 (1.98) | 20.98 (2.01) |
| Height (cm) | 176.81 (8.97) | 172.69 (7.90) | 172.06 (5.84) |
| Weight (kg) | 70.75 (9.64) | 64.25 (10.03) | 64.44 (11.55) |
| Training years (years) | 6.01 (1.49) | 5.58 (0.62) | - |

### Design

We utilized a between-groups 3-way quasi-experimental design. We tested hypotheses regarding differences between athletes and non-athletes, between _interceptive sports_ athletes, _strategic sports_ athletes, and non-athletes, in their scores on the LANT-R task. We grouped participants as athletes or non-athletes based on their self-reported sports participation, where participating in sports once or more per week qualified an individual as an athlete. We grouped participants by sport type based upon Singer’s (2000) article.[2, 27]

### Procedure

The participants were instructed to abstain from alcohol for 24 h and from caffeine-containing substances for 12 h before the experiment. After arriving at the laboratory, the purpose of the study and the instructions for the LANT-R were explained to the participants. After they reported understanding the instructions, the subjects performed the LANT-R task individually in a dimly lit and quiet room. First, they were required to sit in a comfortable chair in front of a computer with their eyes open and were instructed to relax and avoid voluntary movements to minimize motion artifacts. Then, they had to perform a practice block of 24 random trials (described below). Participants were allowed to rest between each block of each trial, and could start the next block once they felt adequately rested. Completing the entire task required approximately 35 min, including the practice and experimental blocks.

### Stimuli

The LANT-R is a simple variation of the revised attention network test [23,28], wherein each stimulus display is rotated 90 degrees clockwise to present lateralized targets. Thus, the stimuli consisted of rows of black arrows pointing upward or downward. The arrows were centered 1.15 degrees to the right or left of the point of eye fixation. A single arrow subtended 0.57 degrees of visual angle, and the contours of adjacent arrows were separated by 0.06 degrees of visual angle. The entire stimulus (target plus flankers) subtended a total 3.09 degrees of visual angle. Each trial began with a fixation cross, and cue boxes were presented in the center of a computer screen during the entire trial. Next, one of the following possible cues for was presented for 100 ms: double cue (both cue boxes flashed), valid cue (one cue box flashed on the location of target stimulus), invalid cue (one cue box flashed in the contralateral location to the target stimulus), or no cue (the stimulus display remained unchanged). Then, there was a fixation period for a variable duration (0, 400, or 800 ms; mean =400 ms). Next, the stimulus was presented and remained for 500 ms. The duration between the onset of the target and the start time of the next trial was one of a set of 12 discrete times from 2,000 to 12,000 ms (10 intervals from 2,000 to 4,250 ms with an increase step of 250 ms, then one 4,750 ms interval and one 12,000 ms interval), approximating an exponential distribution with a mean of 4,000 ms. The mean trial duration was 5,000 ms, with 72 test trails over 4 runs. In each run, all trials were presented in a predetermined counterbalanced order to ensure that each trial type followed every other trial type equally often.

The center arrow was flanked on either side by two arrows in the same direction as the central arrow (congruent condition) or in the opposite direction to that of the central arrow (incongruent condition). Participants were instructed to concentrate on the fixation cross throughout the task. Their task was to identify the direction of the center arrow as fast and accurately as possible by pressing the left key for up-pointing targets and the right key for right pointing targets. Thus, “up” responses were made with the middle finger of the left hand or with the index finger of the right hand, and “down” responses were made with the index finger of the left hand or the middle finger of the right hand. The response hand alternated between blocks in the following order: left, right, left, right. In addition, the response hand alternated between participants in a counterbalanced order. Four blocks were included in this test. Each block contained 48 trials with different cue conditions (valid cue, invalid cue, double cue, or no cue) and flanker conditions (congruent or incongruent). The stimuli were presented and data were recorded using Eprim 2.0 (see Fig. 1).

**Fig. 1.**
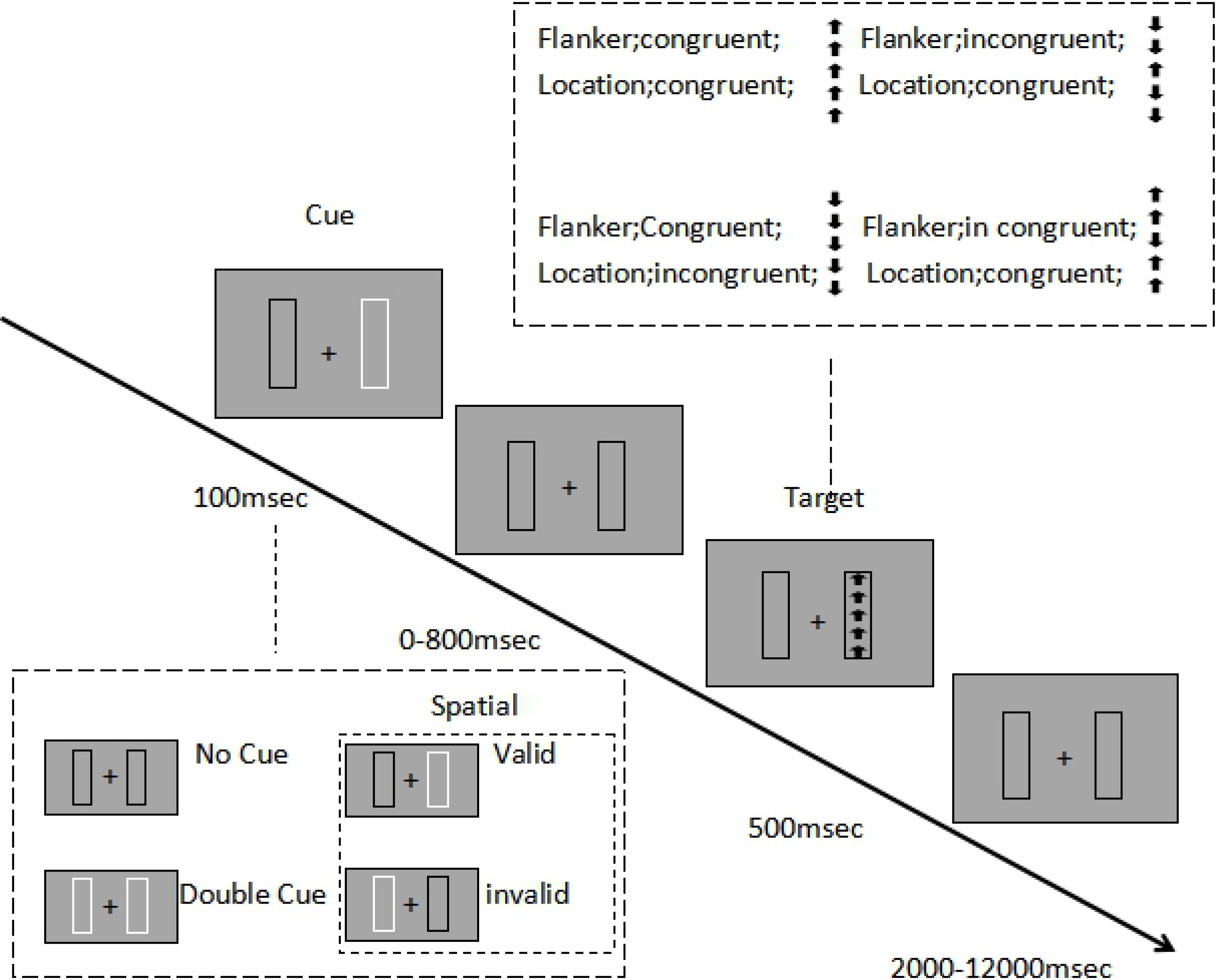
Stimuli and experimental paradigm of the LANT-R

## Measurements

### Reaction time and accuracy

Executive function efficiency of the attentional network was operationally defined based on comparison of RT and ACC performance between congruent and incongruent flanker conditions. We thus calculated a score for each executive function under different cue conditions, as follows.

Flanker conflict effect: (RT/ACC_incongruent_) − (RT/ACC_congruent_)

Flanker conflict effect under no cue condition: (RT/ACC_(no cue, incongruent)_) − (RT/ACC_(no cue, congruent)_)

Flanker conflict effect under double cue condition: (RT/ACC_(double cue, incongruent)_) − (RT/ACC_(double cue, congruent)_)

Flanker conflict effect under valid cue condition: (RT/ACC_(valid cue, incongruent)_) − (RT/ACC_(valid cue, congruent)_)

Flanker conflict effect under invalid cue condition: (RT/ACC_(invalid cue, incongruent)_) − (RT/ACC_(invalid cue, congruent)_)

### fNIRS recording

We used event-related fNIRS to study activation of the executive control network under different cue conditions. Brain activity associated with executive control was determined by subtracting brain activity under the incongruent flanker condition from that under the congruent flanker condition. The spatial cue was thought to add an orienting operation to the central cue prior to the occurrence of the target. The activity associated with different cues was obtained by subtracting the baseline to isolate regions that were more active in response to the spatial cue. We also assessed the changes in blood hemoglobin (Hb) concentration associated with increased neural activity during the LANT-R task using a multi-channel continuous-wave fNIRS instrument (NIRSport, Germany). The fNIRS probe consisted of 8 dual-wavelength sources (760 and 850 nm) and 8 optical detectors, which covered the right FPN (see Fig. 2). The distance between the source and the detector was 3.0 cm. The patch placement was related to the 10-20 system; the location of the 20 channels is shown in Fig. 2. All optodes were arranged on a supporting plastic base and checked for adequate contact on each participant’s scalp. Subsequently, hair under the sensors was brushed away to ensure good skin contact. A channel represented the area measured by one probe-set pair, which was sufficient to measure depths between 2 and 3 cm from the scalp; the channel location was defined as the center position of the pair.

**Fig. 2.**
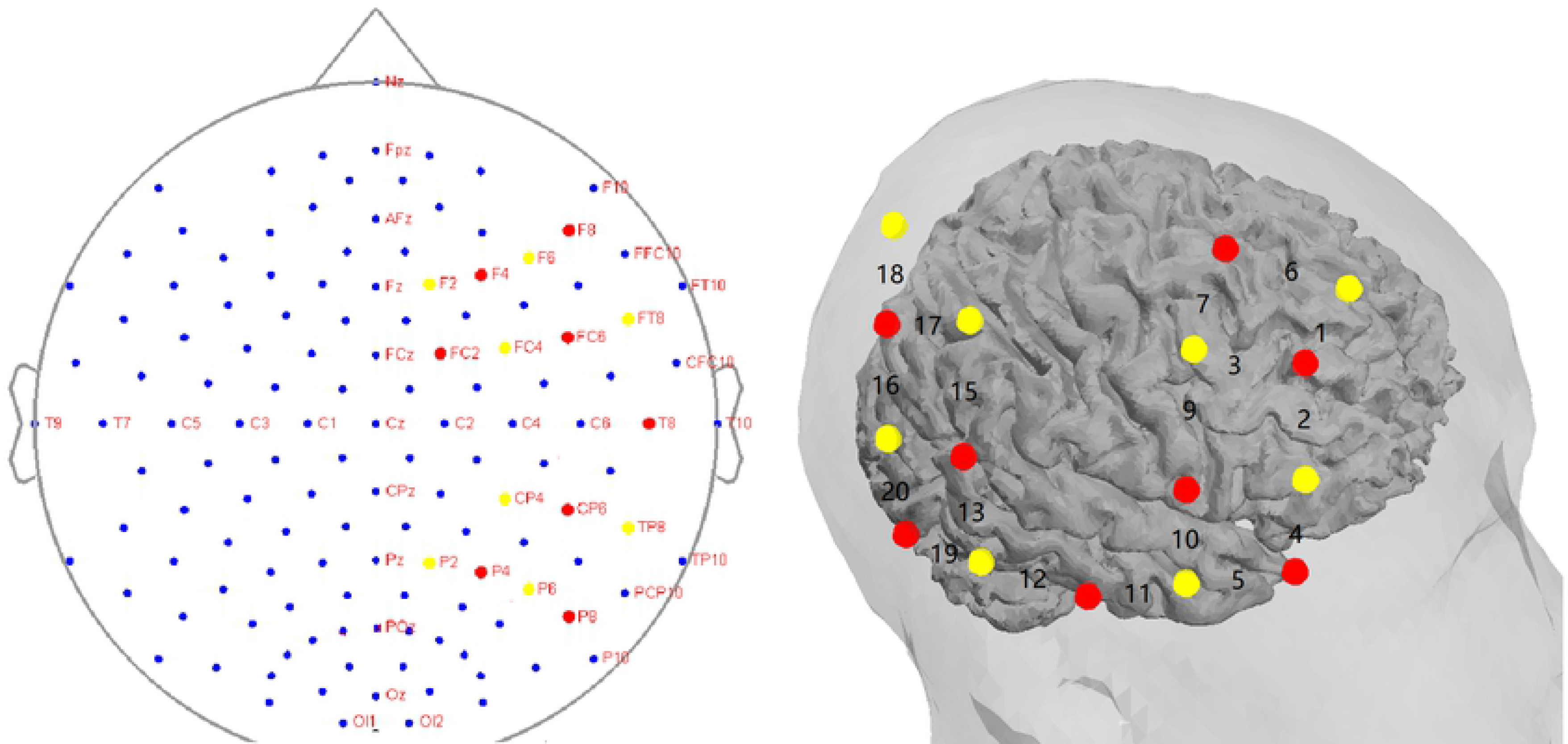
Locations of the sources, detectors, and 20 channels

The sources are represented by the red dots and the detectors are represented by the yellow dots.

## Statistical analyses

### Behavioral data analysis

Statistical analyses of behavioral data were performed using SPSS 21.0 (IBM, New York, USA). The effects of flanker conflict on RT and ACC were analyzed using independent t-tests. Results were considered significant when p < .05. Error trials (incorrect and missing responses) were excluded from the mean RT calculation.

### fNIRS data analysis

The software package Near-Infrared Spectroscopy-Statistical Parametric Mapping (http://bisp.kaist.ac.kr/NIRS-SPM) implemented in the MATLAB environment (MathWorks, Natick, USA) was used to analyze fNIRS data. Gaussian smoothing with a full width at half maximum of 4 s was used to correct for noise. The sampling rate was 7.8 Hz.

Hb data were bandpass filtered between 0.01 Hz and 0.3 Hz to remove baseline drift and physiological noise (e.g., heartbeat). Optical data were converted into Hb signals as arbitrary units using the modified Beer-Lambert Law. General linear model analysis with canonical hemodynamic response curve was performed to analyze oxygenated Hb (HbO) response during experimental conditions. In order to investigate cortical changes in HbO during LANT-R, we selected five regions of interest based on the Brodmann areas (BA) and anatomical locations of the rdLPFC (BA 46; channels 1, 2, and 3), right frontal eye fields (rFEFs; BA 8; channels 6 and 7), right temporoparietal junction (BA19; channels 12, 13, and 19), right posterior parietal cortex (BA 5, 7; channels 16, 17, and 18), and rIFG (BA45, 44; channels 4 and 5). Values for regional changes in HbO were estimated during the task phases from each channel. The values for each region of interest for each participant were acquired in all 288 trials. We recorded the mean concentration change in the HbO signals for 12 s across participants, conditions, and channels. As such, we could compare brain activities of the different participants and analyze the relationship between the activity response and type of athlete. Statistical parametric mapping was performed for group analysis, and HbO was considered significant at an uncorrected threshold of p < 0.05, for stricter analysis.

## Results

### Behavioral data

Overall ACC, overall RTs, and flanker conflict effects on ACC and RT under different cue conditions are presented in Tables 2, 3, 4, and 5, respectively. Analysis revealed that athletes had higher overall ACC, shorter overall RT, and lower flanker conflict effects on RT and ACC compared to non-athlete controls. Further analysis revealed that the flanker effect was due to a stronger conflict effect during the valid cue condition (for ACC and RT) and double cue condition (for RT), but not during the invalid and no cue conditions.

**Table 2.** Effect of conflict on ACC of athletes and non-athletes (*%)*

|  | Overall<br>ACC | Flanker conflict<br>effect | Flanker conflict effect under different cue conditions |  |  |  |
| --- | --- | --- | --- | --- | --- | --- |
|  |  |  | Valid cue | Invalid cue | Double<br>cue | No cue |
| Athletes(n = 32) | 90.10 ± 6.75 | -6.76 ± 6.87 | -4.61 ± 6.03 | -16.07 ± 12.42 | -5.64 ± 10.44 | -9.20 ± 12.46 |
| Non-athletes(n = 24) | 86.33 ± 6.13* | -12.10 ± 9.23* | -11.27 ± 14.50* | -17.90 ± 12.99 | -4.83 ± 13.40 | -11.07 ± 9.45 |
| t | -2.146 | -2.438 | -2.286 | -.527 | .251 | -.627 |
| p | .037 | .018 | .023 | .601 | .803 | .534 |
| d | -0.58 | -0.66 | -0.62 | -0.14 | 0.07 | -0.17 |

Notes: \*P < .05, \*\*P < .01. ACC = accuracy

**Table 3.** Effect of conflict on RT of athletes and non-athletes (*ms)*

Notes: \*P < .05, \*\*P < .01. RT = reaction time
|  | Overall RT | Flanker conflict<br>effect | Flanker conflict effect under different cue conditions |  |  |  |
| --- | --- | --- | --- | --- | --- | --- |
|  |  |  | Valid cue | Invalid cue | Double cue | No cue |
| Athletes(n = 32) | 576.10 ± 67.60 | 84.81 ± 22.33 | 63.58 ± 24.18 | 120.63 ± 49.32 | 97.00 ± 35.62 | 113.53 ± 44.12 |
| Non-athletes(n = 24) | 624.80 ± 50.96** | 101.60 ± 31.09* | 86.17 ± 22.65** | 126.43 ± 42.63 | 119.83 ± 38.02* | 105.83 ± 44.77 |
| t | 2.944 | 2.227 | 3.152 | -.456 | 2.255 | -.632 |
| p | .005 | .032 | .003 | .650 | .029 | .530 |
| d | 0.79 | 0.60 | 0.85 | -0.12 | 0.61 | -0.17 |

**Table 4.**
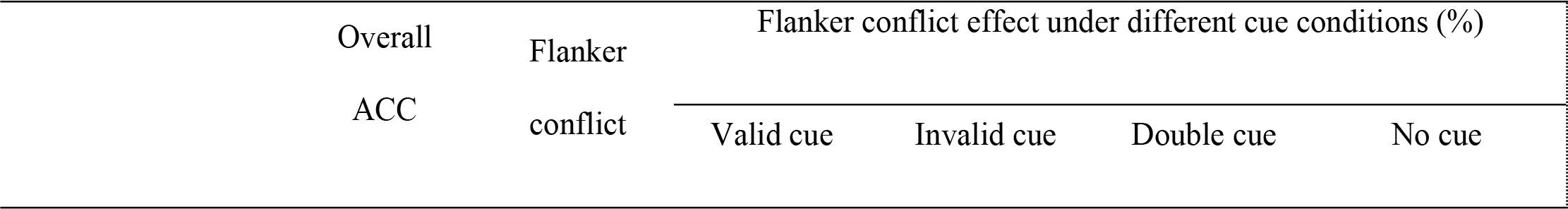
Effects of conflict on ACC of athletes: interceptive vs. strategic sports (*%)*

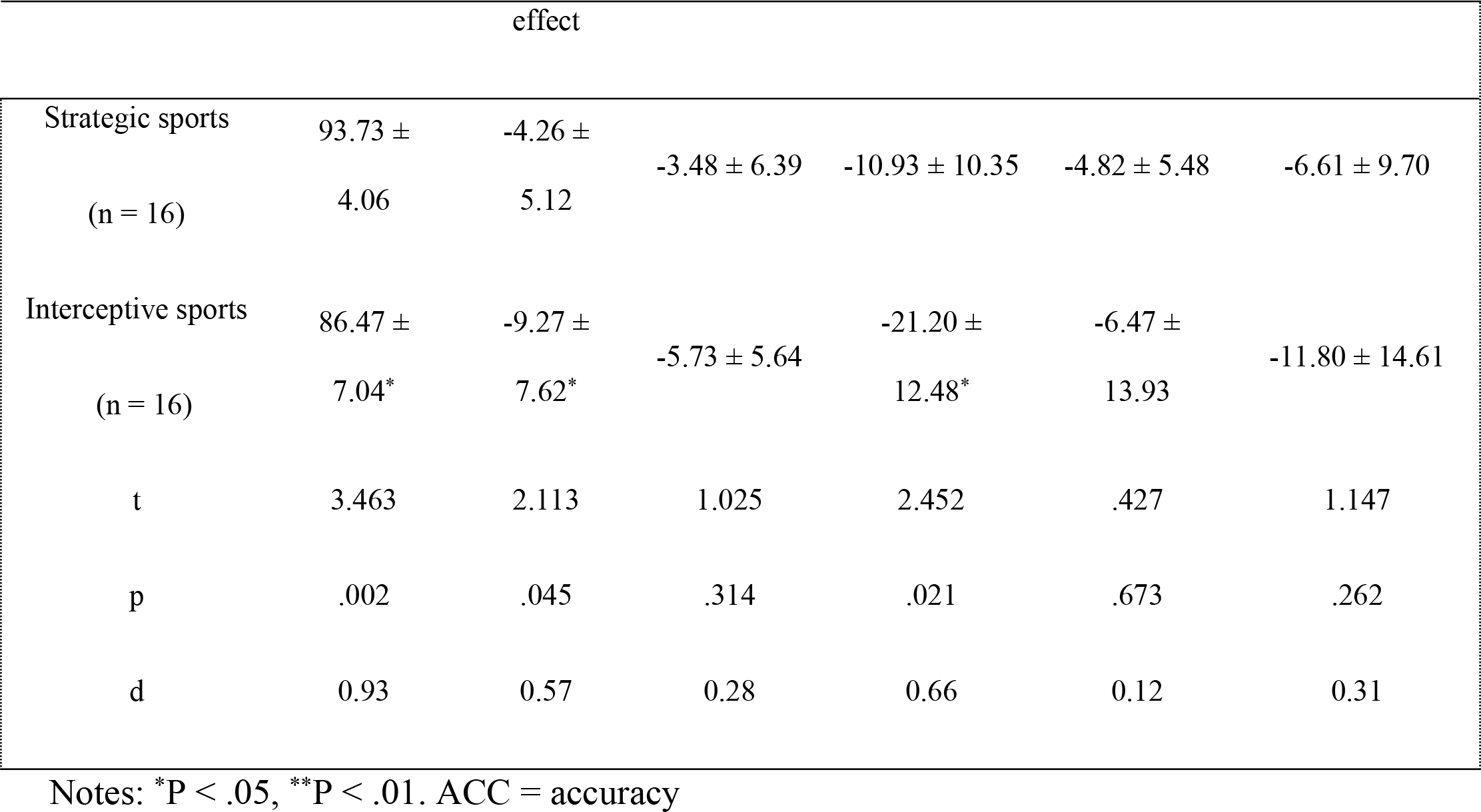

**Table 5.** Effects of conflict on RT of athletes: interceptive vs. strategic sports (*ms)*

|  | Overall RT | Flanker<br>conflict effect | Flanker conflict effect under different cue conditions (ms) |  |  |  |
| --- | --- | --- | --- | --- | --- | --- |
|  |  |  | Valid cue | Invalid cue | Double cue | No cue |
| Strategic sports (n = 16) | 618.56 ± 58.95 | 94.26 ± 19.22 | 72.83 ± 24.16 | 149.53 ± 35.27 | 90.07 ± 31.21 | 119.80 ± 48.42 |
| Interceptive sports (n = 16) | 533.64 ± 46.14** | 75.35 ± 21.71* | 54.33 ± 21.08* | 103.33 ± 37.11** | 103.93 ± 39.39 | 107.27 ± 40.03 |
| t | 4.393 | 2.525 | 2.234 | 3.495 | -1.069 | .773 |
| p | .000 | .018 | .034 | .002 | .295 | .446 |
| d | 1.18 | 0.68 | 0.60 | 0.94 | -0.29 | 0.21 |

Notes: \*P<.05, \*\*P<.01. RT = reaction time.

Comparison of strategic and interceptive sports athletes revealed that strategic athletes had higher ACC and lower flanker conflict effects on ACC, as well as longer RT and greater flanker conflict effects on RT. Furthermore, the flanker effect was due to a stronger conflict effect during the valid cue condition (for ACC and RT) and the invalid cue condition (on RT), but not during the double or no cue conditions.

### fNIRS data

The cortical activation associated with the flanker conflict effect had contrasting effects during the LANT-R between the athlete and non-athlete groups. Activation in the rdLPFC was generally greater in the athletes than in controls (Fig. 3a). Moreover, we detected stronger activation in the right posterior parietal cortex (Fig.3b) when participants in the strategic sport group detected and resolved conflict.

**Fig. 3.**
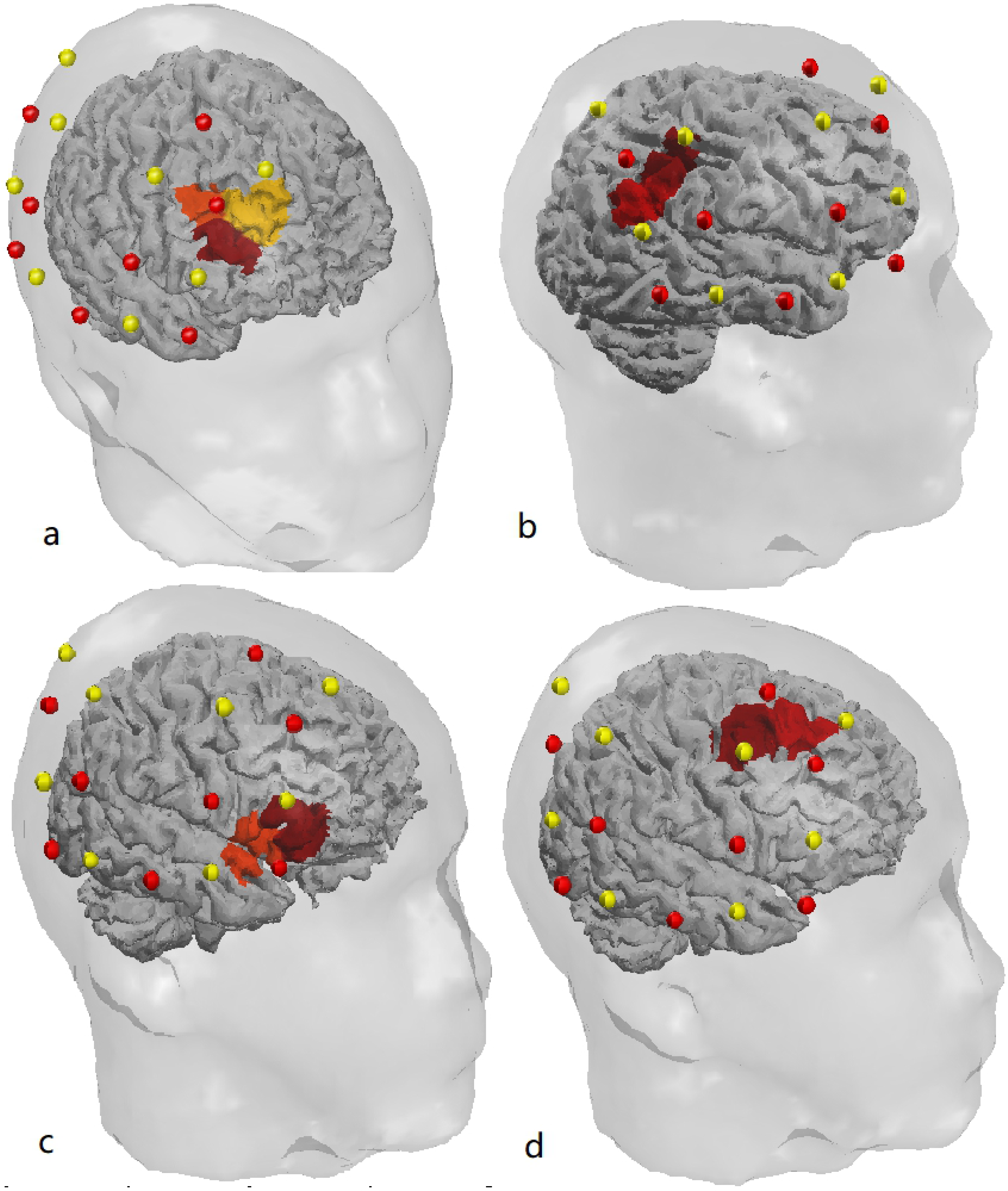
Activation associated with flanker conflict and cue type. a. Athletes vs non-athletes with flanker conflict b. Strategic vs interceptive with flanker conflict c. Strategic vs interceptive under valid and invalid cue conditions d. Athletes vs non-athletes under valid and double cue conditions

### Activation associated with cue type

Table 6 presents data on the differences between athletes and non-athletes encountering different cue stimulations. Athletes had more significant activation in the rFEF (Fig. 3d) under the valid cue and double cue conditions. Furthermore, we compared brain activation following valid cue and invalid cue stimulation in the strategic sports group to that in the interceptive sports group to determine whether there were differences in the activation of the IFG (Fig. 3c).

## Discussion

The present study evaluated the differences of the neural mechanism use of LANT-R to identify the characteristics of executive control in athletes from open-skill sports using fNIRS. The results indicated superiority of the athlete group in processing flanker conflicts; in addition, there were differences in processing strategy between strategic and interceptive sports athletes which strategic sports involve more distinct TD processes. These effects could be attributed to different activity in the right frontoparietal network.

As for the behavioral results, we found substantial support for our hypotheses. First, as expected, athletes exhibited a shorter RT and higher ACC compared to non-athletes. These findings were consistent with those of previous studies suggesting that athletes have better reaction speed and validity than non-athletes and novices on cognitive tests [13]. We speculate that these findings can be explained in part by highly efficient cognitive processing among athletes, which contributes to speed and accuracy of decision-making during game play. Alternatively, the superior cognitive performance may be due to the cognitive benefits of physical activity [31].

In addition, athletes revealed significantly smaller conflict effects on RT and ACC relative to non-athletes. It is likely that athletes have a better ability to perceive and solve conflicts (i.e., higher executive control efficiency) [32]. Another possibility is that athletes may be able to more efficiently allocate attention, an important determinant of success in sports such as basketball, badminton, or volleyball that involve rapid changes in visual information [33]. The *most intriguing* finding of the current study is that valid cue and double cue exert a positive influence on executive control in athletes. That means that orienting and alerting facilitates conflict processing. This result is consistent with those reported by Fan et al. [23]. To the best of our knowledge, this study is the first to demonstrate a facilitatory effect of alerting on executive control. One might speculate that athletes tend to recruit more brain regions directly related to executive control for resolving conflict, taking advantage of cues more efficiently[34]. Athletes have a distinguished ability to maintain higher cognitive resource commitment and more efficient resource utilization, and some, for example table tennis athletes, have higher discrimination ability for spatial vision [15].

Second, we compared the flanker effects on RT and ACC between the interceptive and strategic sport groups. As expected, athletes in the strategic group exhibited slower overall RT and larger conflict effects on RT (but not on ACC). Further analysis revealed that this conflict effect is due to the stronger flanker incongruency under the invalid cue condition. In brief, strategic sports athletes detected and resolved flanker conflicts in more time than interceptive sports athletes while achieving a better ACC, particularly following an invalid cue condition. This suggests that strategic sport athletes do not focus attention on the target location in advance, as they did not move their attention to the precise location of the upcoming target under the invalid spatial cue conditions. As a result, they had more time to resolve the flanker conflict while under time pressure to achieve the desired outcome, resulting in higher ACC.

In contrast, interceptive athletes displayed faster RT and smaller flanker effects on RT, but not higher ACC. These results suggest that interceptive athletes’ ability to focus effectively on a target is inferior to strategic sports athletes. This may owe to variations in the flanker effect due to the characteristics of each type of sport. Parallel top-down (TD) and bottom-up (BU) processes occur in the attention network during motor learning, training, and competition. It is necessary to integrate these processes based on the skill set required for each sport. The key integration process in interceptive sports comprises reducing TD processes to exploit sensory feedback (BU) in order to automatically select and guide motor actions within limited time periods. This may explain why interceptive athletes displayed superior RTs. In contrast, strategic sports involve maximizing TD processes under time pressure in order to observe and anticipate, by way of example, a ball’s trajectory, an opponent’s position and intention, and a teammate’s position. The athlete then deploys the tactic according to past experience or the coach’s arrangement and subsequently uses his or her body to manipulate the movements of the above objects. It is likely that the increased flanker effect on RT provided time for these processes, which are required for anticipation and/or tactical layout. In sports, these abilities are called “game intelligence” or “Olympic brain”.

Furthermore, we used fNIRS analysis to confirm the above-mentioned neural processing strategy. As shown in S1 Table, athletes displayed greater activity in the rdLPFC than non-athletes while detecting and resolving flanker conflicts. Activation of the rdLPFC is an important part of TD processing in the right frontoparietal network [24]. This further validates athletes’ higher resource commitment during executive control processes, which results in enhanced neural activation of the rdLPFC. The coupling of enhanced cognitive performance and increased activation in the prefrontal cortex has been suggested by Byun and Yanagisawa [7, 32]. In other words, the training enhanced the athletes’ ability to recruit brain resources for the executive control task. Additionally, from the fNIRS results, we observed more recruitment in rDLPC and rIFG in detecting and resolving conflict during TD processing in the basketball group compared with the other groups. This was accompanied by higher accuracy of synchronous behavioral performance (i.e., incongruent condition-congruent condition). Hence, we speculate that the high accuracy of basketball athletes, who are involved in strategic sports, might be attributed to higher activation in rDLPFC and rIFG. This would indicate that more recruitment during executive control processing guaranteed efficiency of information processing under conflict conditions. The coactivation of rIFG and rdLPFC during cognitive processing supported the hypothesis that basketball athletes’ have a strategic advantage in TD processing. In a “speed-accuracy trade-off”, they adopted preference towards accuracy, rather than reaction speed.

We also investigated activation under different cue conditions between athletes and non-athletes, and between interceptive sports athletes and strategic sports athletes. We found markedly greater activation in the rFEF in the athlete group during the valid and double cue conditions, further supporting a facilitatory effect of orienting and alerting on executive control. Functional anatomical studies indicate that the frontal eye field mediates the covert allocation of attention to visual locations across a wide variety of visual tasks [24, 30, 33].

Another important finding of the present study was that activity in the rIFG involving action observation and anticipation [8] was higher in strategic sports athletes than in interceptive sports athletes during the invalid and double cue conditions. These findings were consistent with our hypothesis and the behavioral results, suggesting that superior anticipation ability in strategic sports athletes is correlated with higher cortical activity [5, 34].

Our study has numerous limitations. These limitations, as listed below along with suggestions for how they may be avoided in future research, weaken the statistical power of our study to some extent. Although our sample size (n = 48) was large enough to yield significant results and large effect sizes, we suggest future research to use a larger sample. Because our study was quasi-experimental, we were unable to determine causality. We suggest using a pre-test/post-test design, in which participants are randomly assigned to strategic sports athlete, interceptive sports athlete, and non-athlete groups. If executive functions improve through athletic training and competition in a manner consistent with our findings, this would be strong evidence that certain types of sports differentially improve these functions.

## Conclusions

The current study provides neural evidence substantiating the hypothesis that open skill athletes demonstrate significantly more efficient executive control when compared to non-athletes. This was accompanied by increased activity in the rdLPFC. Moreover, they had a more remarkable ability to utilize cues. Athletes from interceptive sports demonstrated increased speed when solving conflict, while those from strategic sports demonstrated higher accuracy. In addition, differences in attentional processing were found between interceptive sports athletes and strategic sports athletes facing similar cues. In particular, following an invalid cue, TD control appears to play an important role when strategic sports athletes must make a cautious decision. This can be attributed to the right frontoparietal network.

## Acknowledgements

We thank all the participants for their participation. This work was supported by a project from Ministry of Science and Technology of the People’s Republic of China (grant number 2018YFF0300401).

